# Evidence that lower socioeconomic position accentuates genetic susceptibility to obesity

**DOI:** 10.1101/074054

**Authors:** Jessica Tyrrell, Andrew R Wood, Ryan M Ames, Hanieh Yaghootkar, Robin N. Beaumont, Samuel E. Jones, Marcus A Tuke, Katherine S. Ruth, Rachel M Freathy, George Davey Smith, Stéphane Joost, Idris Guessous, Anna Murray, David P. Strachan, Zoltán Kutalik, Michael N Weedon, Timothy M Frayling

**Author notes:** Corresponding Author: Timothy Frayling Corresponding Author Address: Genetics of Complex Traits, Institute of Biomedical and Clinical Science, University of Exeter Medical School, Royal Devon and Exeter Hospital, Barrack Road, Exeter, EX2 5DW, UK.

## Abstract

Susceptibility to obesity in today’s environment has a strong genetic component. Lower socioeconomic position (SEP) is associated with a higher risk of obesity but it is not known if it accentuates genetic susceptibility to obesity. We aimed to use up to 120,000 individuals from the UK Biobank study to test the hypothesis that measures of socioeconomic position accentuate genetic susceptibility to obesity. We used the Townsend deprivation index (TDI) as the main measure of socioeconomic position, and a 69-variant genetic risk score (GRS) as a measure of genetic susceptibility to obesity. We also tested the hypothesis that interactions between BMI genetics and socioeconomic position would result in evidence of interaction with individual measures of the obesogenic environment and behaviours that correlate strongly with socioeconomic position, even if they have no obesogenic role. These measures included self-reported TV watching, diet and physical activity, and an objective measure of activity derived from accelerometers. We performed several negative control tests, including a simulated environment correlated with BMI but not TDI, and sun protection use. We found evidence of gene-environment interactions with TDI (P_interaction_=3×10^−10^) such that, within the group of 50% living in the most relatively deprived situations, carrying 10 additional BMI-raising alleles was associated with approximately 3.8 kg extra weight in someone 1.73m tall. In contrast, within the group of 50% living in the least deprivation, carrying 10 additional BMI-raising alleles was associated with approximately 2.9 kg extra weight. We also observed evidence of interaction between sun protection use and BMI genetics, suggesting that residual confounding may result in evidence of non-causal interactions. Our findings provide evidence that relative social deprivation best captures aspects of the obesogenic environment that accentuate the genetic predisposition to obesity in the UK.

## INTRODUCTION

The prevalence of obesity is set to dramatically exceed targets set by the World Health Organisation and place an increasingly large burden on health services throughout the world^1^. Whilst environmental influences, including diet and lifestyle have caused the obesity epidemic^2^, twin and family studies show that genetic factors influence susceptibility to obesity in today’s environment^3,4^. Recent genetic studies have identified many common genetic variants associated with BMI ^5^ but the role of genetic susceptibility in different modern day environments has proven controversial.

Different studies have concluded that physical inactivity^6,7^, consuming more fried food^8^, more fizzy drinks^9^ or more protein^10^ accentuate the risk of obesity in those genetically predisposed. These studies have been limited by the necessity of using large meta-analyses of studies with heterogeneous measures of the environment or have been hindered by the potential biases caused by the use of self-reported measures of the environment. Some of these studies have claimed evidence of causation for the environmental factor tested, with such statements as “these data support a causal relationship among the consumption of sugar-sweetened beverages, weight gain and the risk of obesity”; however unlike main effect Mendelian randomisation studies, gene x environment studies are susceptible to confounding^11^. A recent study, testing only the *FTO* locus, overcame many of these issues by using a single large, relatively homogeneous study, the UK Biobank, and testing many measures of the environment in the same statistical model^12^.

Social deprivation is correlated with obesity in children^13^ and adults^14^. There is evidence that individuals from more deprived backgrounds make poorer food choices^15^ and tend to be less active^16^. Whilst individuals from more socially deprived backgrounds are more overweight on average, few studies have tested the hypothesis that deprivation accentuates genetic susceptibility to obesity, with the exception of the recent study using the UK Biobank that found nominal evidence that deprivation accentuates the effect of the variant at the *FTO* locus^12^. We used a single large, relatively homogeneous study, the UK Biobank, to test this hypothesis. We provide evidence that relative social deprivation in the UK accentuates genetic predisposition to obesity. We also suggest that socioeconomic position and residual confounding may explain some of the previously described interactions between BMI genetics and specific aspects of the obesogenic environment.

## RESULTS

### Social deprivation is associated with BMI and variance in BMI in the UK Biobank study

Social deprivation, as measured by the Townsend deprivation index (TDI) was associated with BMI in UK Biobank individuals – both in the whole study (Supplementary table 1) and the subset of 119,464 with genetic data available at the time of our analysis (Table 1).

**Table 1:** Demographics of the 119,733 individuals in the UK Biobank with Townsend deprivation index, BMI and genetic data available by deprivation.

|  | Most deprived | Least deprived | P* |
| --- | --- | --- | --- |
| N | 59,861 | 59,872 |  |
| Mean age at recruitment (SD) | 56.5 (8.1) | 57.4 (7.8) | <1x10<sup>-15</sup> |
| Male, N (%) | 28,306 (47.3) | 28,383 (47.4) | 0.91 |
| Mean Townsend deprivation index (SD) | 0.83 (2.5) | -3.78 (0.9) | <1x10<sup>-15</sup> |
| Mean BMI (SD) | 27.9 (5.1) | 27.2 (4.5) | <1x10<sup>-15</sup> |
| Obese, N (%) | 16,647 (27.9) | 13,233 (22.1) | <1x10<sup>-15</sup> |
| Current smoker, N (%) | 9,193 (15.4) | 4,139 (6.9) | <1x10<sup>-15</sup> |
| Type 2 diabetes, N (%) | 2,304 (3.9) | 1,702 (2.8) | <1x10<sup>-15</sup> |
| Coronary artery disease, N (%) | 3,312 (5.5) | 2,449 (4.1) | <1x10<sup>-15</sup> |
| High job class*, N (%) | 25,947 (43.4) | 29,540 (49.3) | <1x10<sup>-15</sup> |
| Mean education duration (SD) | 14.2 (5.3) | 15.2 (4.9) | <1x10<sup>-15</sup> |

\* High job class included the following job categories: associate professionals, professional occupations and managers and senior officials.

Individuals who were more socially deprived had higher BMIs and a higher prevalence of obesity. A one standard deviation (1SD) higher Townsend deprivation index was associated with a 0.42 kgm^−2^ [95%CI: 0.40, 0.45] higher BMI (*p* = <1×10^−15^). The 50% of individuals who were most deprived (n=59,861) had an average BMI of 27.9 kgm^−2^, and 95% had a BMI between 21.0 and 37.4 kgm^−2^ (a range of 16.4) whereas the 50% who were least deprived (n=59,872) had an average BMI of 27.2 kgm^−2^, and 95% had a BMI between 21.1 and 35.4 (a range of 14.3).

### A BMI genetic risk score is associated with BMI in the UK Biobank study

The BMI genetic risk score, consisting of 69 known BMI-associated variants, was associated with higher BMI and explained 1.5% of the variation in BMI, a figure consistent with previous studies^5^.

### A higher level of deprivation is associated with an accentuated genetic susceptibility to higher BMI

The effect of the BMI genetic risk score on BMI was larger in the group of 50% living in the most relatively deprived situations (0.025 standard deviations per allele [95%CI: 0.023-0.027]) compared to the group of 50% living in the least deprived situations (0.022 SDs per allele [95%CI: 0.020-0.024]) (Table 2; Figure 1A). When performing the analysis with Townsend deprivation index on a continuous scale (a more robust analysis than using dichotomized measures) the evidence of interaction was strong: *P*_*interaction*_6×10^−12^ (*P*_*interaction*_ 2×10^−10^, using robust standard errors). This apparent gene x deprivation interaction meant that, compared to below average deprivation (in the UK Biobank), above average deprivation was associated with a 0.92kgm^−2^ higher BMI in people with the highest genetic risk (top decile) but a 0.35kgm^−2^ higher BMI in people at least genetic risk (bottom decile)(Table 2, Figure 1A). Another way of expressing the interaction is that, within the 50% group living in the most deprived situations, carrying 10 additional BMI-raising alleles (weighted by effect size) was associated with approximately 3.8 kg extra weight in someone 1.73 metres tall. In contrast, within the 50% group living in the least deprived situations, carrying 10 additional BMI-raising alleles was associated with approximately 2.9 kg extra weight in someone 1.73 metres tall. These differences were even stronger when using a cut off that reflected the UK population average TDI (Supplementary table 2). Because UK Biobank participants are of higher socioeconomic position (SEP) than the average person of this generation in the UK, we also used a standard deviation value of -0.4 TDI to define above and below average deprivation. This UK Biobank value approximates to the UK population mean based on number of people with an undergraduate degree (33% versus 20%^17^). The BMI genetics – TDI interaction effects between the 29.6% of UK Biobank individuals with the lowest TDI and the 70.4% with the highest TDI were even stronger.

**Figure 1:**
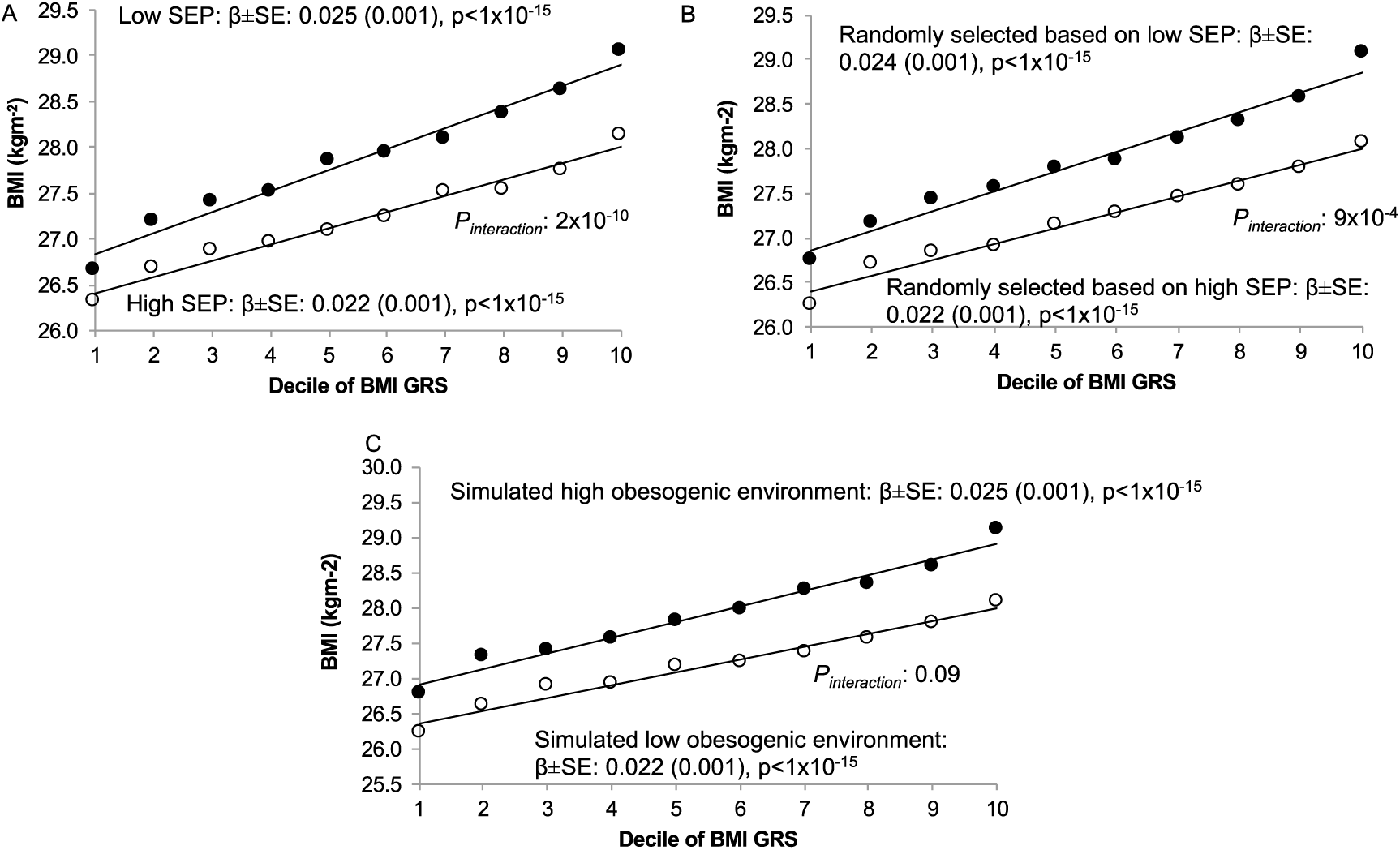
Association between the BMI genetic risk score (by decile) and BMI in A) most socially deprived (black circles) and least socially deprived (white circles); B) individuals randomly selected to be of high BMI (black circles) and individuals randomly selected to be of low BMI (white circles) and C) individuals in the high obesogenic simulated environment (black circles) and individuals in the low obesogenic simulated environment (white circles). Note for the simulated environment we used the median BMI GRS BMI association after 1000 simulations. For B it was not possible to use a continuous measure in the calculation of the interaction term. This figure is based on a similar way of showing interaction data with a BMI genetic risk score from ^33^ SEP: Socioeconomic Position

**Table 2:** Differences in BMI by BMI genetic risk score decile (kgm^2^) and by allele (inverse normalised scale) for a) actual Townsend deprivation index, b) randomly selected groups based on the BMI distribution in the two TDI categories and b) simulated environments with a similar association to TDI and BMI. There are slightly fewer individuals in the randomly selected groups due to the need to obtain an identical distribution to the real TDI variable.

| Trait | Obesogenic category | N | BMI (SD) | BMI difference in 10% lowest genetic risk | BMI difference in 10% highest genetic risk | Per-allele Beta | SE | P association | P interaction* | P Interaction Robust** |
| --- | --- | --- | --- | --- | --- | --- | --- | --- | --- | --- |
| Townsend Deprivation Index (natural scale) | High SEP<br>TDI≤-2.294 | 59,872 | 27.20<br>(4.47) | +0.35 $\text{kgm}^{-2}$ | +0.92 $\text{kgm}^{-2}$ | 0.022 | 0.001 | $<1 \times 10^{-15}$ | **$6 \times 10^{-12}$** | **$2 \times 10^{-10}$** |
| | Low SEP<br>TDI$>$-2.294 | 59,861 | 27.87<br>(5.13) | | | 0.025 | 0.001 | $<1 \times 10^{-15}$ | | |
| Townsend Deprivation Index (natural scale)*** | High SEP<br>TDI≤-2.294 | 42,753 | 27.09<br>(4.32) | +0.35 $\text{kgm}^{-2}$ | +0.90 $\text{kgm}^{-2}$ | 0.022 | 0.001 | $<1 \times 10^{-15}$ | **$1 \times 10^{-7}$** | **$1 \times 10^{-6}$** |
| | Low SEP<br>TDI$>$-2.294 | 42,684 | 27.72<br>(4.94) | | | 0.024 | 0.001 | $<1 \times 10^{-15}$ | | |
| Townsend Deprivation Index (Inverse normalised) | High SEP<br>TDI≤-2.294 | 59,872 | 27.20<br>(4.47) | +0.35 $\text{kgm}^{-2}$ | +0.92 $\text{kgm}^{-2}$ | 0.022 | 0.001 | $<1 \times 10^{-15}$ | **$7 \times 10^{-4}$** | **$8 \times 10^{-4}$** |
| | Low SEP<br>TDI$>$-2.294 | 59,861 | 27.87<br>(5.13) | | | 0.025 | 0.001 | $<1 \times 10^{-15}$ | | |
| Randomly selected individuals (TDI on natural scale)**** | Pseudo low risk | 59,753 | 27.19<br>(4.47) | +0.51 $\text{kgm}^{-2}$ | +1.00 $\text{kgm}^{-2}$ | 0.022 | 0.001 | $<1 \times 10^{-15}$ | **$8 \times 10^{-4}$** | **$9 \times 10^{-4}$** |
| | Pseudo high risk | 59,711 | 27.86<br>(5.12) | | | 0.024 | 0.001 | $<1 \times 10^{-15}$ | | |
| Simulated environment | Pseudo low risk | 59,741 | 27.16<br>(4.61) | +0.58 $\text{kgm}^{-2}$ | +1.00 $\text{kgm}^{-2}$ | 0.022 | 0.001 | $<1 \times 10^{-15}$ | **0.093** | **0.094** |
| | Pseudo high risk | 59,740 | 27.90<br>(5.01) | | | 0.025 | 0.001 | $<1 \times 10^{-15}$ | | |
| Sun protection use | Usually or always use | 68,507 | 27.32<br>(4.75) | +0.32 $\text{kgm}^{-2}$ | +0.63 $\text{kgm}^{-2}$ | 0.022 | 0.001 | $<1 \times 10^{-15}$ | **$1 \times 10^{-4}$** | **$1 \times 10^{-4}$** |
| | Never or sometimes use | 50,561 | 27.81<br>(4.89) | | | 0.025 | 0.001 | $<1 \times 10^{-15}$ | | |

BMI adjusted for age, sex, ancestral principal components and assessment centre location. Models additionally adjusted for genotyping platform
\* Interaction p-value
\*\* Interaction p-value accounting for heteroscedasticity using robust standard errors
\*\*\*Interaction model adjusted for TV watching, physical activity and Western diet.
\*\*\*\*by Meta-heuristic sampling

Previous studies have suggested a stronger gradient by genetic risk for BMI at younger rather than later ages ^18^. We stratified our analyses by age and saw consistent interaction effects for TDI across different age groups (Supplementary table 3). We also noted that the interaction effect was not driven by specific BMI associated variants, but was a feature of general genetic susceptibility to higher BMI, as measured by the 69 SNP BMI risk score (Supplementary table 4 & supplementary figure 1). Excluding the FTO variant did not alter the evidence of interaction.

In the CoLaus study of 5,237 individuals from Switzerland, we did not observe any evidence that TDI interacted with the BMI genetic risk score, but the effect estimates overlap those in the UK Biobank (Supplementary table 5).

#### Lower occupational job class was not associated with an accentuated genetic susceptibility to higher BMI

In both the UK Biobank and the 1958 Birth Cohort individuals with lower job classes had a higher mean and standard deviation for BMI. However, there was no evidence of an interaction between job class and genetic risk score in determining BMI in either study (Supplementary table 5).

#### Less time spent in education was not associated with an accentuated genetic susceptibility to higher BMI

In the UK Biobank individuals spending less time in education had a higher mean and standard deviation for BMI. However, there was no evidence of an interaction between time in education and genetic risk score in determining BMI (Supplementary table 5).

#### Are previously described interactions between BMI genetics and specific measures of the environment driven by socioeconomic position?

We next hypothesized that the interaction between genetic susceptibility to higher BMI and socioeconomic position (as measured by TDI) may result in evidence of interactions with specific measures of the environment correlated with TDI. Previous studies have observed stronger effects of BMI raising alleles in groups of individuals who are less active^6,19^, eating more fried food^8^ and consuming more sugary drinks^9^. However, all of these groups were more overweight on average than those with the healthier lifestyles and any interaction observed may have been a feature of higher BMI and lower socioeconomic position, not the specific environment tested. We tested 12 additional measures of the environment, including an objective measure of activity based on accelerometer wear. One of the measures was a negative control, sun protection use in the summer, which has no clear obesogenic role, but is strongly correlated with TDI (supplementary methods). Five of these 12 variables showed evidence of interaction at p<0.05 and including TDI as a covariate did not appreciably alter the evidence of interaction (Supplementary table 6). Our negative control, using less sun protection in the summer, was associated with higher deprivation and showed evidence of interaction with genetic susceptibility to higher BMI, before and after adjustment for TDI (Table 2).

#### Testing the specificity of the TDI interaction using a simulated environment and groups of individuals randomly selected to be of different BMIs

We next performed two additional analyses to test the specificity of the interaction to TDI. These tests represented “impossible by the proposed mechanism” negative controls. ^20^ These analyses also helped limit the possibility that statistical artefacts were influencing our results, such as wider variances in BMI in deprived individuals compared to less deprived individuals.

First, we used a heuristic approach to randomly select groups of individuals with identical BMI means and distributions to the high and low TDI groups and tested for evidence of interaction. This was repeated 100 times. The associations between the BMI genetic risk score and BMI in these randomly selected individuals were similar to those observed when we selected based on Townsend deprivation index, but none of the 100 analyses showed an interaction p-value lower than the real TDI interaction (median p=9×10^−4^, Table 2, Figure 1B, Figure 2A). We note that this analysis is not completely representative of the real TDI analysis because the interaction term is a binary variable (presence or absence of the individual in the randomly selected groups of higher and lower BMI) not continuous.

**Figure 2:**
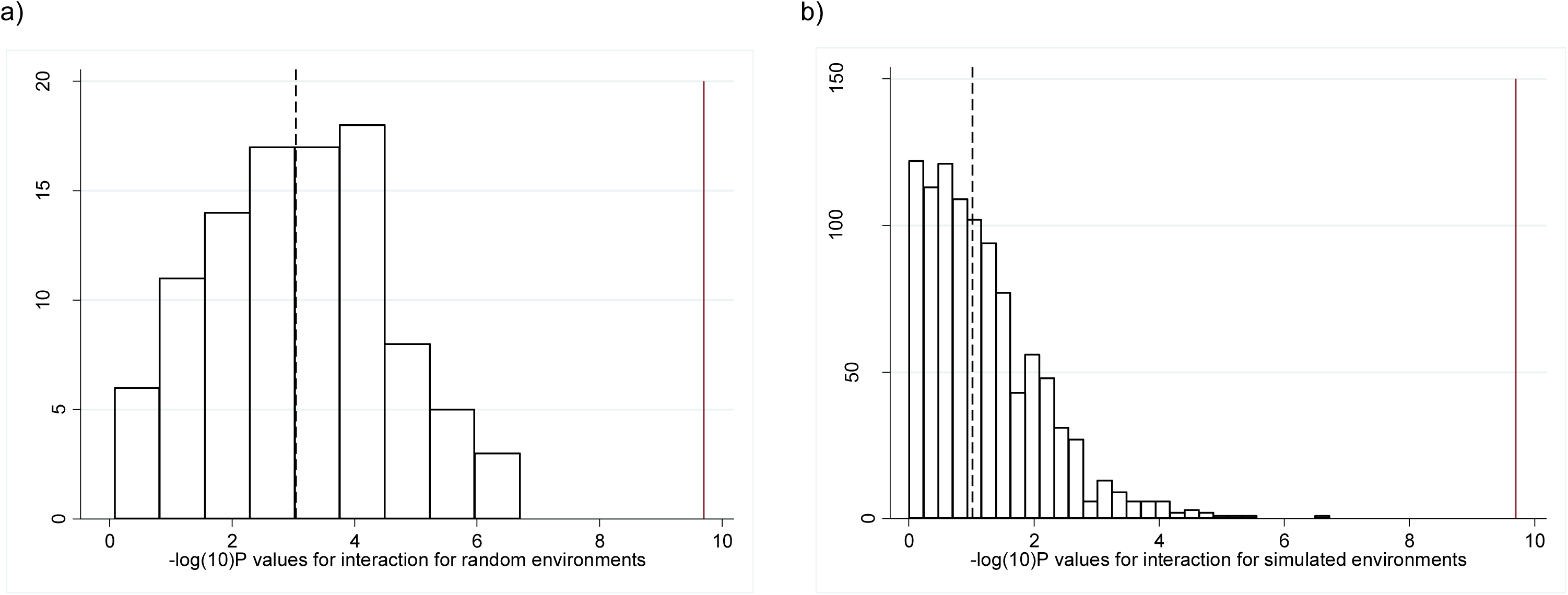
Histogram showing the −log_10_(P values) for the interactions from a) the meta-heuristic generated environment and b) the simulated environments. The dashed line represents the median value and the solid red line represents the p -value obtained from the real data.

Second, we generated a continuous variable as a dummy environment by regressing TDI against BMI, the BMI GRS, age, sex and age squared. The fitted values were then added to random permutations of the residuals to generate dummy environments that were associated with BMI to the same extent as TDI but not associated with TDI. We repeated this 1000 times and each time tested for evidence of interaction using the dummy continuous environment as an interaction term. When dichotomized into two equal groups, the BMI means, distributions and BMI genetic risk score associations with BMI were similar to those observed in the two groups defined by the real Townsend deprivation index (Table 2). However, no evidence for interaction was observed (p=0.09 based on median of 1000 dummy environments, Figure 1C & Figure 2B). In the 1000 permutations of a dummy environment, we never observed evidence as strong as that observed with real TDI, providing evidence at p<0.001 that the TDI effect was genuine (Figure 2B).

#### Sensitivity analyses

The evidence of interaction between TDI and the BMI genetic risk score remained when we adjusted for other measures of the environment and behaviour previously associated with gene-obesity interactions *P*_*interaction*_ 9×10^−8^ (*P*_*interaction*_ 1×10^−6^, using robust standard errors). These factors included self-reported physical activity, TV watching, and a more “westernized” diet (reflecting a more calorie dense diet – see methods). All of these measures were associated with TDI in the UK Biobank (Supplementary table 7). These factors may be on the causal pathway from relative deprivation to higher BMI, but the fact that no one measure accounts for the TDI-BMI genetic risk score association suggests that any interaction is not driven by one particular aspect of the environment associated with TDI. We addressed several other factors that could affect our results in the supplementary information.

## DISCUSSION

In the UK Biobank we found evidence that relative deprivation, as measured by the Townsend Deprivation Index, accentuates genetic susceptibility to higher BMI. The corollary of our findings is that exposure to lower levels of relative deprivation may partially attenuate the effects of genetic susceptibility to obesity. Our results are important because they provide further evidence for public health campaigns aimed at reducing obesity but suggest that measures that target more deprived individuals may have proportionally higher impact. The mechanism by which higher levels of deprivation may accentuate the genetic risk of obesity is not known. When adjusting for measures of physical activity, a more calorie dense “westernised” diet and sedentary activity, the evidence of interaction remained strong. This observation suggests that no one aspect of the obesogenic environment we tested can explain the interaction effect with TDI, although a caveat to this argument is that these other measures were self-reported. The evidence of interaction remained strong when adjusting for urban versus rural dwelling, an objective measure associated with obesity in the UK Biobank and previously proposed as a contributory factor to the obesogenic environment (through reduced exposure to open spaces for example^21^). A further implication of our results is that they suggest that no one aspect of the obesogenic environment we tested has a specific role in accentuating the genetic susceptibility to obesity. This conclusion contrasts with that from some previous studies reporting interactions with TV watching, physical activity, fried food and sugary drink consumption^6-10^. Including sun protection use as a negative control helped us control for the possibility that TDI does not capture all of the variance in socioeconomic position. The importance of using negative controls in epidemiology to control for this residual confounding has been discussed^20^ and is closely related to the use of one of Hill’s original criteria for causal inference in epidemiology – that of specificity of the exposure-outcome association^22^. The fact that this negative control showed significant interaction, even after adjustment for TDI, suggests that either it is a bad negative control or it is correlated with other obesogenic factors not captured by TDI. Low vitamin D levels (which would be caused by high use of sun protection) are associated with higher BMI, but there is genetic evidence that this is not a causal relationship^23^, and even if it were, would have resulted in evidence of interaction in the opposite direction to our observation. Further work, including the use of negative controls that are likely associated with unmeasured confounders but are implausible, will help disentangle which aspects of the environment are causally interacting with BMI genetics to accentuate the risk of high BMI.

Our results are consistent with data from twins, where there is evidence that the genetic component to obesity is stronger in young UK children exposed to the modern environment (twins born in the 1990s and measured at the age of 9), compared to measures from twin studies in earlier generations^3^ and that the genetic and environmental components to common traits varies by UK region ^24^.

Although the interaction effect sizes observed were small based on each allele, when summed across all alleles, the interaction effects we observed were meaningful – for example, in the group of 50% in the most deprived situations, carrying 10 additional BMI-raising alleles was associated with 3.8 kg extra weight in someone 1.73m tall. In contrast, in the group of 50% in the least deprived situations, carrying 10 additional BMI-raising alleles was associated with 2.9 kg extra weight in someone 1.73m tall. The BMI genetic risk score is associated predominantly with extra fat mass with a smaller effect on lean mass, and 900 grams of extra fat per person will equate to a large effect on obesity related disease in the relatively deprived population. Furthermore, participants in the UK Biobank are less deprived than average UK citizens which may mean the effects are larger in the general population.

Our analysis had a number of strengths. The major strength was the availability of a single large study, which was beneficial for two main reasons. First, it provided us with relatively homogenous measures of the environment. Several previous studies were limited to meta-analyses of summary statistics from many studies with heterogeneous measures of the environment although the most recent also used the UK Biobank and used the individual level data to jointly model multiple exposures and provide evidence that some measures we did not test, including frequency of alcohol consumption, remain significant when adjusting for TDI ^12^.

Second, it allowed us to test the robustness of our results by randomly selecting individuals and testing interactions using a dummy, simulated environment – an important negative control experiment to control for potential statistical artefacts and non-specific interaction effects. A third advantage is that we used an objective measure of the environment, which provides a cleaner interpretation of results compared to those from previous studies that have had to rely on subjective measures such as self-reported diet and physical activity. These subjective measures are often complex mixtures of environment and behaviour and may be subject to reporting biases. The fourth advantage of our study is that we used a negative control variable, sun protection use, which helps control for residual confounding.

There were a number of limitations to this study. First, no evidence of interaction between TDI and BMI genetics was noted in a smaller study of 5,237 individuals from the CoLaus Study. However, the power of this analysis was limited by the small sample size. Additionally CoLaus was from an urban based study in another country (Switzerland) and as such might not be comparable to the UK Biobank. No evidence of interaction was noted for job class or number of years in education. These variables were available in <70% of the individuals and are only weakly correlated with TDI (r=0.15 and 0.11 respectively) and it is possible that TDI captures the key components of the obesogenic environment in those living in relatively deprived situations better than these variables. The lack of interaction with job class could possibly be explained by manual workers doing more exercise, as observed in the UK Biobank. Secondly, we cannot rule out the possibility that statistical artefacts have at least partially influenced our results. We observed some evidence of interaction when randomly sampling individuals and using dummy environments, although the evidence was much weaker and some residual evidence of interaction may be expected given that the dummy environments are still slightly correlated with TDI. When the outcome (here BMI) and the interaction term (here TDI) are correlated, and when the variance of the outcome and interaction term are correlated, interaction analyses are prone to artefacts. However, we performed a number of sensitivity analyses that suggested these artefacts were not responsible for the interaction, including testing both BMI and TDI on different scales. Previous studies of BMI gene x environment interactions have not necessarily accounted for such sources of potential error. Thirdly, the UK Biobank is enriched for people of lower levels of deprivation compared to the average in the UK, but this bias would be expected to accentuate the effect of genetic variants in people of low deprivation, the opposite to what we observed. It is also important to note that we were not testing for specific gene variant-environment interactions but instead asking a question of public health relevance – are people at higher risk of obesity due to their genetics more susceptible to the obesogenic environment? It may be that particular genetic variants, for example those acting through genes affecting appetite, may have different effects compared to others, for example those acting through metabolic rate. Finally, variation in BMI in today’s environment is estimated to be 40% genetic (although there are wide confidence intervals around all estimates) but our genetic risk score only accounts for 1.5% variance in BMI. However, there is no reason to think that the 69 genetic variants used do not provide a general snapshot of general genetic susceptibility to high BMI.

In conclusion, analyses in 119,733 individuals from the UK Biobank provides evidence that socioeconomic position, as measured by the Townsend deprivation index, best captures the aspects of the environment that accentuate the genetic risk of obesity.

## METHODS

### UK Biobank participants

The UK Biobank recruited over 500,000 individuals aged 37-73 years (99.5% were between 40 and 69 years) in 2006-2010 from across the UK. Participants provided a range of information via questionnaires and interviews (e.g. demographics, health status, lifestyle) and anthropometric measurements, blood pressure readings, blood, urine and saliva samples were taken for future analysis: this has been described in more detail elsewhere^25^. We used up to 119,688 individuals of British descent from the initial UK Biobank with BMI available. We did not include other ethnic groups, because individually they were underpowered to detect previously reported effects. British-descent was defined as individuals who both self-identified as white British and were confirmed as ancestrally Caucasian using principal components analyses (PCA) of genome wide genetic information. This dataset underwent extensive central quality control (http://biobank.ctsu.ox.ac.uk) including the exclusion of the majority of third degree or closer relatives from a genetic kinship analysis of 96% of individuals. We performed an additional round of principal components analysis (PCA) on the UK Biobank participants of British descent. We selected 95,535 independent single nucleotide polymorphisms (SNPs) (pairwise *r*^2^ <0.1) directly genotyped with a minor allele frequency (MAF) ≥ 2.5% and missingness <1.5% across all UK Biobank participants with genetic data available at the time of this study (*n*=152,732), and with HWE *P*>1×10^−6^ within the white British participants. Principal components were subsequently generated using FlashPCA ^26^ and the first five adjusted for in all analyses.

### Patient Involvement

Details of patient and public involvement in the UK Biobank are available online (http://www.ukbiobank.ac.uk/about-biobank-uk/ and https://www.ukbiobank.ac.uk/wp-content/uploads/2011/07/Summary-EGF-consultation.pdf?phpMyAdmin=trmKQlYdjjnQIgJ%2CfAzikMhEnx6).

### Phenotypes

#### BMI

The UK Biobank measured weight and height in all participants and calculated BMI. BMI was available for 119,464 individuals of white descent with Townsend deprivation indices and genetic data available. We performed analyses of BMI on both its natural (kgm^−2^) and an inverse normalised scale to account for differences in variances (see below).

#### Obesogenic environment

##### Townsend deprivation index

The Townsend deprivation index is a composite measure of deprivation based on unemployment, non-car ownership, non-home ownership and household overcrowding; a negative value represents high socioeconomic position ^27^. This was calculated prior to joining the UK Biobank and is based on the preceding national census data, with each participant assigned a score corresponding to the postcode of their home dwelling. For presentation purposes, the Townsend deprivation index variable was dichotomised at the median with 59,872 individuals in the least deprived group (Townsend deprivation index <-2.294) and 59,861 in the most deprived group (Townsend deprivation index >-2.294).

The Townsend deprivation index variable was skewed (Supplementary figure 2) and therefore we single inverse normalised this variable for use in sensitivity analyses.

##### Job class

The UK Biobank asked people to select their current or most recent job. This was classified into one of the following strata: elementary occupations, process plant and machine operatives, sales and customer service occupations, leisure & other personal service occupations, personal service occupations, skilled trades, admin and secretarial roles, business and public sector associate professionals, associate professionals, professional occupations and managers and senior officials. Data were available for 76,374 individuals. The job class variable was dichotomised at the median value with 38,942 individuals in the high job class category (associate professionals and above) and 37,374 in the low job class category.

##### Years in education

A variable based on the standardised 1997 International Standard Classification of Education (ISCED) of the United Nations Educational, Scientific and Cultural Organisation was created in the UK Biobank, using previously published guidelines ^28^. Data were available for 118,775 individuals. The educational years variable was dichotomised into shorter educational duration (7-15 years, n=63,572), versus a longer educational duration (19 to 20 years, n=55,203).

##### Specific measures of the obesogenic environment

To test the hypothesis that previously published BMI gene-environment interactions may have been driven by SEP we tested 12 additional measures of the obesogenic environment, including self-reported and objectively measured physical activity, several diet variables, TV-watching and, as an implausible control variable, sun protection use (Supplementary table 8).

### Replication with TDI: CoLaus Study

The CoLaus Study ^29^ is a population based study including over 6500 participants from Lausanne (Switzerland). This study included inhabitants aged 35-75 years at baseline (2003-2006) and they were followed up between 2009 and 2012 (mean follow-up 5.5 years). Within this cohort TDI was available for 5,237 individuals with BMI and BMI genetic variants available. The use of TDI in Lausanne may capture socioeconomic position in a different way to the UK Biobank, because, for example, not owning a car correlates with higher SEP. The CoLaus study complied with Declaration of Helsinki and was approved by the local Institutional Ethics Committee.

### Replication with job class: 1958 Birth Cohort

The 1958 Birth Cohort ^30^ has followed persons born in England, Scotland and Wales during one week in 1958 from birth into middle age. Within this cohort 6,171 individuals had information on social class based on their own current or most recent occupation (at age 42), body mass index (measured at age 44-45) and genetic data. The social class measure was dichotomised separately for men and for women, yielding 2,873 participants in the high job class category and 3,298 in the low job class category.

### Selection of Genetic Variants associated with BMI and Genetic Risk Score

We selected 69 of 76 common genetic variants that were associated with BMI at genome wide significance in the GIANT consortium in studies of up to 339,224 individuals (Supplementary table 9)^5^. We used these variants to create a genetic risk score to represent genetic susceptibility to high BMI – we were not testing specific variants for interaction, but instead how genetic susceptibility overall may be influenced by environmental and behavioural exposures. We used genotypes imputed by central UK Biobank analysts using a combined reference panel of the 1000Genomes and UK10K sequenced datasets. We confirmed the validity of the genotypes by testing the genetic risk score for association with BMI. We limited the BMI SNPs to those that were associated with BMI in the analysis of all European ancestry individuals and did not include those that only reached genome-wide levels of statistical confidence in one-sex only, or one-strata only. Variants were also excluded if known to be classified as a secondary signal within a locus. Three variants were excluded from the score due to potential pleiotropy (rs11030104 [*BDNF* reward phenotypes], rs13107325 [*SLC39A8* lipids, blood pressure], rs3888190 [*SH2B1* multiple traits]), 3 SNPs not in Hardy Weinberg Equilibrium (*P*<1×10^−6^; rs17001654, rs2075650, rs9925964) or the SNP was unavailable (rs2033529).

The imputed dosages for each SNP were recoded to represent the number of BMI-increasing alleles for that particular SNP. A BMI genetic risk score (GRS) was created using the SNPs. Each allele associated with high BMI was weighted by its relative effect size (β-coefficient) obtained from the previously reported BMI meta-analysis data^5^. A weighted score was created (equation 1) in which β is the β-coefficient representing the association between each SNP and BMI.

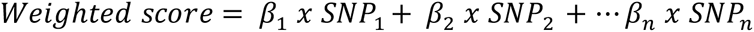

The weighted score was rescaled to reflect the number of BMI-increasing alleles (Equation 2).

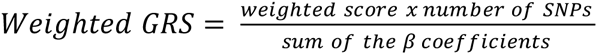

### Statistical analysis

The mean and standard deviation of BMI was calculated for two groups in the UK Biobank: those living in the 50% relatively most and least deprived situations.

For each of the measures of the socioeconomic position we calculated the association between the 69 SNP BMI GRS and BMI in the high risk and low risk environments using linear regression models. BMI was adjusted for age, sex, five ancestry principal components and assessment centre location. We additionally adjusted for genotyping platform (two were used) at runtime.

Interactions between the genetic variables and the obesogenic environment variables on BMI were tested by including the respective interaction terms in the models (e.g. interaction term= GRS x Townsend deprivation index (continuous)). A continuous measure for Townsend deprivation index was used to limit spurious results from the gene x environment interactions.

We performed the analyses in two ways. First we analysed the data with BMI on its natural scale (kgm^−2^) (residualised for age, sex, centre location and five ancestry principal components). Second we inverse normalised the data so that BMI, in each strata had a mean BMI of 0 and a SD of 1. This analysis allowed us to account for the differences in BMI variation observed in high and low risk strata. We present primary results from the inverse normalised data. To further assess the extent to which differences in BMI variation could influence our results we tested for heteroscedasticity using the Breusch-Pagan test as implemented with the estat hettest in STATA^31^. Standard regression analysis can produce biased standard errors if heteroscedasticity is present ^32^. If heteroscedasticity was present we utilised robust standard errors, using the vce(robust) option in STATA, which relaxes the assumption that errors are both independent and identically distributed and are therefore more robust.

We also repeated the analysis adjusting for other measures of the environment previously associated with interactions, including self-reported physical activity, TV watching and diet^6,7,9,10^.

The analysis was repeated using single inverse normalised Townsend deprivation index.

Finally, we investigated each of the 69 SNPs individually. The association between each of the 69 SNPs and BMI in the high risk and low risk environments was investigated using linear regression models. Interactions between each SNP and the Townsend deprivation index on BMI were tested by including the respective interaction terms in the models (e.g. interaction term= SNP x Townsend deprivation index (continuous)).

Identical analyses were performed in the CoLaus Study and the 1958 Birth Cohort.

### Randomly selecting groups of individuals to be of different BMIs

Firstly, we employed a meta-heuristic sampling approach to randomly select 2 groups of individuals with BMI distributions identical to the high and low Townsend deprivation index groups. This method selected 59,712 individuals with a mean BMI of 27.86 and a standard deviation of 5.12 and a group of 59,754 individuals with a mean BMI of 27.19 and a standard deviation of 4.47. There was no overlap between individuals selected for the two groups. This analysis was repeated 100 times. No significant overlap between the groups was observed (i.e. those in the high risk group in 1 analysis were randomised across high and low risk groups in subsequent analyses).

### BMI genetic risk score interactions with dummy “environments”

Secondly, we created a random dummy continuous environmental variable that was correlated with BMI to approximately the same extent as the Townsend deprivation index (r^2^=0.0894), but was only minimally correlated with Townsend deprivation index itself. The new variable, Y, was created in STATA by regressing TDI on BMI, the genetic risk score and a range of covariates (age, age^2^, sex) and taking the fitted values and the residuals. The residuals were then randomly permuted 1000 times and added to the fitted values. This ensures that the simulated variable has the same conditional expectations and same residual distributions as the real TDI variable. Further information on this method is provided in the supplement. The interaction model was run for all 1000 simulations.

### Negative control variable – self reported sun protection use

We used sun protection use as a negative control variable to attempt to account for residual confounding. UK Biobank participants were asked “Do you wear sun protection (e.g. sunscreen lotion, hat) when you spend time outdoors in the summer?" with the options: Never, Sometimes, Most of the time, Always, Don’t go out in the sun, Don’t know and Prefer not to answer. The variables was correlated with TDI and BMI but is implausible as a mechanism (see discussion for why vitamin D exposure is unlikely to be a mechanism in this context).

## Acknowledgements

This research has been conducted using the UK Biobank Resource.

## Funding Information

J.T. is funded by a Diabetes Research and Wellness Foundation Fellowship. S.E.J. is funded by the Medical Research Council (grant: MR/M005070/1) M.A.T., M.N.W. and A.M. are supported by the Wellcome Trust Institutional Strategic Support Award (WT097835MF). A.R.W. H.Y. and T.M.F. are supported by the European Research Council grant: 323195:SZ-245 50371- GLUCOSEGENES-FP7-IDEAS-ERC. R.M.F. is a Sir Henry Dale Fellow (Wellcome Trust and Royal Society grant: 104150/Z/14/Z). R.B. is funded by the Wellcome Trust and Royal Society grant: 104150/Z/14/Z. R.M.A is supported by the Wellcome Trust Institutional Strategic Support Award (WT105618MA). Z.K. is funded by Swiss National Science Foundation (31003A-143914). The funders had no influence on study design, data collection and analysis, decision to publish, or preparation of the manuscript. The CoLaus study was supported by research grants from GlaxoSmithKline, the Faculty of Biology and Medicine of Lausanne, Switzerland, and the Swiss National Science Foundation (grants 33CSCO-122661, 33CS30-139468 and 33CS30-148401.

## Author Contributions

J.T., T.M.F., M.N.W. designed the study. J.T., H.Y., T.M.F.,M.N.W. wrote the manuscript. S.E.J.,J.T., R.B., K.S.R., A.R.W., M.A.T., H.Y., R.M.F., A.M., M.N.W., S.J., Z.K. performed data processing, statistical analyses and interpretation. T.M.F is the guarantor. T.M.F affirms that the manuscript is an honest, accurate, and transparent account of the study being reported; that no important aspects of the study have been omitted; and that any discrepancies from the study as planned (and, if relevant, registered) have been explained.

## Conflict of Interest

All authors have completed the ICMJE uniform disclosure form at http://www.icmje.org/coi_disclosure.pdf and declare: no support from any organisation for the submitted work; M.N.W has received speakers fees from Ipsen and Merck, and T.M.F has consulted for Boehringer Ingelheim, no other relationships or activities that could appear to have influenced the submitted work.

## Data access

The data reported in this paper are available via application directly to the UK Biobank.

